# Mapping of Facial and Vocal Processing in Common Marmosets with ultra-high field fMRI

**DOI:** 10.1101/2023.08.09.552659

**Authors:** Audrey Dureux, Alessandro Zanini, Stefan Everling

**Affiliations:** Centre for Functional and Metabolic Mapping, Robarts Research Institute, University of Western Ontario, London, ON N6A 5K8, Canada; Department of Physiology and Pharmacology, University of Western Ontario, London, ON N6A 5K8, Canada

**Author notes:** Corresponding author: Audrey Dureux, Centre for Functional and Metabolic Mapping, Robarts Research Institute, University of Western Ontario, London, Canada.

**Keywords:** Faces, vocalizations, audiovisual, multisensory, awake marmosets, fMRI, social communication

## Abstract

Primate communication relies on multimodal cues, such as vision and audition, to facilitate the exchange of intentions, enable social interactions, avoid predators, and foster group cohesion during daily activities. Understanding the integration of facial and vocal signals is pivotal to comprehend social interaction. In this study, we acquired whole-brain ultra-high field (9.4T) fMRI data from awake marmosets (*Callithrix jacchus*) to explore brain responses to unimodal and combined facial and vocal stimuli. Our findings reveal that the multisensory condition not only intensified activations in the occipito-temporal ‘face patches’ and auditory ‘voice patches’ but also engaged a more extensive network that included additional parietal, prefrontal and cingulate areas, compared to the summed responses of the unimodal conditions. By uncovering the neural network underlying multisensory audiovisual integration in marmosets, this study highlights the efficiency and adaptability of the marmoset brain in processing facial and vocal social signals, providing significant insights into primate social communication.

## Introduction

Primates emit a variety of signals during daily social communication, expressing specific emotional states, intentions, activities, or responses to external environmental features ^1^. These complex signals encompass visual, tactile, olfactory and auditory cues ^2^, with facial expressions and vocalizations serving as the primary sources for face-to-face primate communication, a notion already postulated by Charles Darwin ^3^. The crossmodal integration of facial expressions and vocalizations is vital for perceiving a conspecific’s vocalization and concurrent facial behavior ^4,5^. Functional magnetic resonance imaging (fMRI) studies have revealed that humans, Old-World macaque monkeys, and New-World marmosets share a face processing system, consisting of interconnected patches distributed across the temporal and prefrontal cortex. In humans, face-selective patches are found in the lateral occipital cortex, the fusiform gyrus and in anterior and posterior regions of the superior temporal sulcus (STS) ^6–8^. In macaques, these patches are located along the occipitotemporal axis mainly along the STS and in the frontal cortex ^9–15^. In marmosets, similar patches have been identified along occipitotemporal axis and in lateral frontal cortex ^16–18^, following a similar organization as in macaques and humans ^11,19^.

Several studies have also associated the processing of negative facial expressions with higher activations in temporal face-selective regions and in prefrontal and subcortical areas in both humans and macaques ^6,20–25^, a pattern that we also recently observed in marmosets ^18^. Furthermore, vocalizations and vocal production contribute to interaction and cohesion within primate groups ^26–28^. fMRI studies in humans have identified three voice-selective patches located along the mid-superior temporal gyrus to the anterior superior temporal gyrus (TVAa, TVAm, TVAp) and in premotor and inferior frontal areas ^29–35^.

In macaques, two clusters in the STS have been identified with stronger activations for vocalizations than for other sounds categories ^36–39^. Additionally, the recent discovery of a vocalization-selective cluster in the macaque anterior temporal pole suggests a similar functional organization of higher-level auditory cortex in macaques and humans ^39^. These results suggest that vocalization processing is organized in ‘voice patches’ in the temporal lobe, analogous to the well-establish ‘face-patches’ ^36,40,41^.

Recently, we identified in marmosets a network akin to what was seen in humans. Vocalization-selective activations were observed in temporal, frontal and anterior cingulate cortices ^42^. Furthermore, three voice patches were discerned along the STS, potentially homologous to the three human voice patches ^42,43^.

The association between facial expressions and specific vocalizations enables a nuanced understanding of social cues, enhancing communication and social bonding among primate groups ^5^. Yet, even though facial and vocal patches in primates have been individually studied, the process of integrating these multisensory signals during social interactions remains intricate and not fully understood.

Recent studies have started to uncover the neural substrates of multisensory integration in primates, demonstrating that audiovisual integration occurs in specific regions of both the monkey face-patch and voice-patch systems ^44–47^ as well as in the ventral lateral prefrontal cortex (VLPFC) ^48–50^. Human studies have highlighted the involvement of temporal ^51,52^, frontal ^53,54^, and parietal ^53^ cortices in processing combined visual and auditory information.

The present study aims to extend the understanding of multisensory integration in primates by examining the specific neural circuits that enable the combined processing of facial and vocal cues in the common marmoset (*Callithrix jacchus*). Utilizing ultra-high field MRI, we acquired whole-brain fMRI data from six awake marmosets while the animals were presented with videos of conspecific faces with no sound, conspecific vocalizations with no video, videos of conspecific faces with corresponding vocalizations, and scrambled versions of each of these conditions.

Our objective was to identify the networks involved in the processing of face and vocal signals, as well as their integration, to contribute to a more comprehensive understanding of marmoset social communication. Our results provide a novel perspective on the neural architecture of audiovisual multisensory integration for social stimuli in marmosets.

## Results

Our primary objective was to uncover the neural architecture responsible for processing and integrating face and vocal signals in marmosets. Specifically, we sought to identify the brain regions responsive to face and vocalization processing, and the regions that are activated by these combined signals. To this end, we employed a sparse fMRI block-design, presenting six different conditions to awake marmosets: a unimodal condition of marmoset face videos, a unimodal condition of vocalization sounds, a multimodal condition of videos of marmoset faces combined with corresponding vocalizations, and the respective scrambled versions of these conditions. Our analysis included the creation of conjunction maps to explore both specific and common activations among these conditions, and the examination of the superadditive effect to investigate the responses to combined audiovisual stimulation.

### Functional brain activations during the processing of visual signals

Initially, we examined the processing of marmoset face videos and their corresponding scrambled versions, compared to a baseline period where only a central dot was presented on the screen. The group activation maps for each condition are depicted in Figures 1A and 1D. Marmoset face videos (Figure 1A) recruited a bilateral network primarily along the occipitotemporal axis, encompassing visual areas V1, V2, V3, V4, V4T, MT, V6, dorsointermediate part (19DI), lateral and inferior temporal areas TE1, TE2, TE3, TEO, the fundal area of the superior temporal sulcus (FST), PGa-IPa, and ventral temporal areas 35, 36. The scrambled face videos (Figure 1D) activated visual areas V1, V2, V3, V4T, MT, and the FST area. Subcortically, bilateral pulvinar, lateral geniculate nucleus (LGN) and right amygdala were recruited by face videos, whereas no activations were observed for scrambled faces.

**Figure 1.**
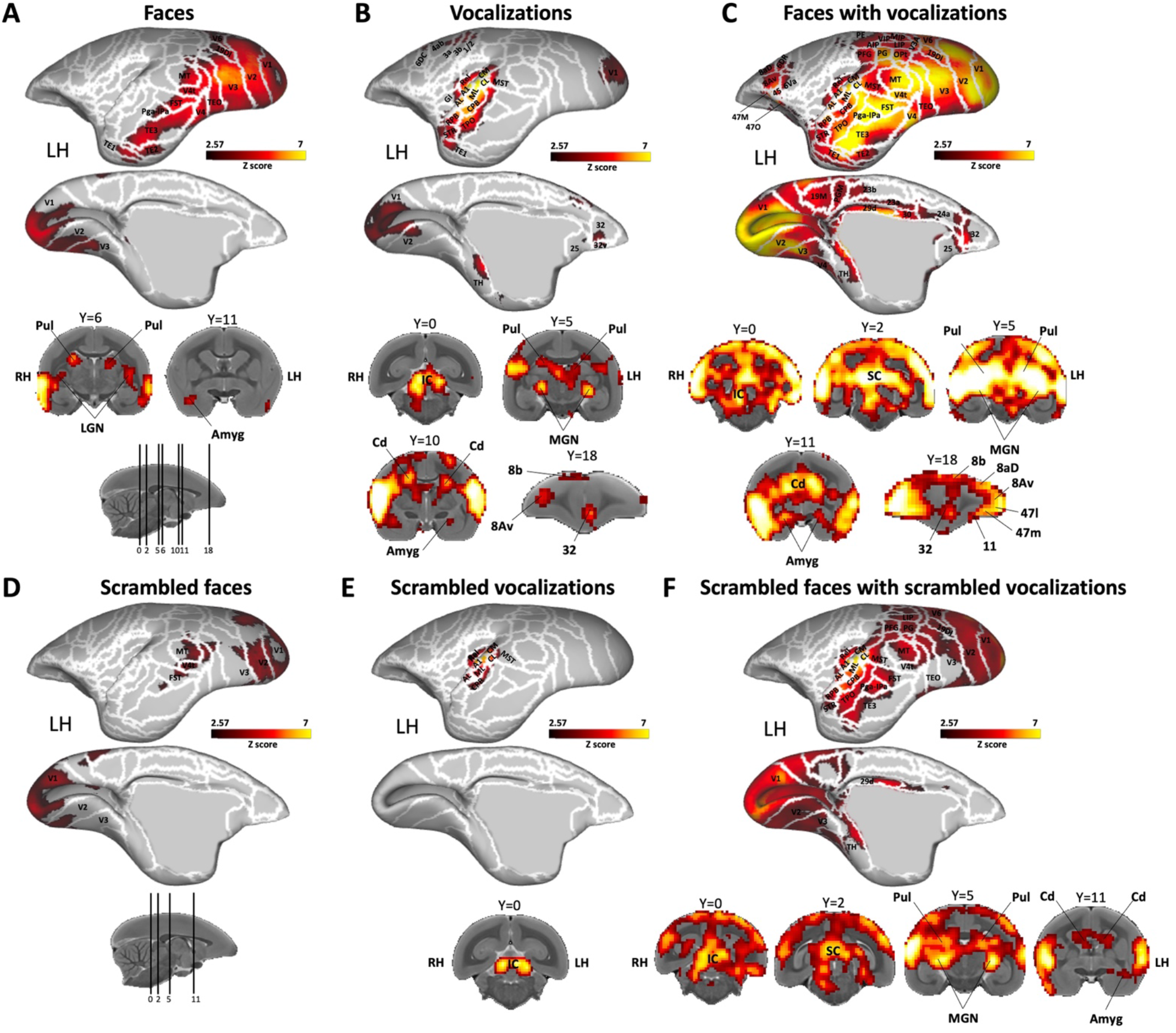
Brain networks activated by each condition *versus* baseline. This figure represents the group functional map for each condition, showing significantly greater activations compared to baseline: marmoset face videos (A), marmoset vocalizations (B), marmoset face videos with corresponding vocalizations (C), scrambled marmoset face videos (D), scrambled marmoset vocalizations (E), and scrambled marmoset face videos with corresponding scrambled vocalizations (F). These group maps are based on data from six awake marmosets and are displayed on both lateral and medial views of the fiducial marmoset cortical surfaces, left hemisphere. Subcortical activations are represented on coronal slices. The white line delineates the regions based on the Paxinos parcellation of the NIH marmoset brain atlas ^86^. The reported brain areas meet an activation threshold corresponding to z-scores > 2.57 (p<0.01, AFNI’s 3dttest++, cluster-size correction α=0.05 from 10000 Monte-Carlo simulations).

### Functional brain activations during the processing of auditory signals

Next, we explored the activation patterns for vocalization processing by analyzing each auditory condition – marmoset vocalizations and scrambled marmoset vocalizations – compared to the baseline period. Figures 1B and 1E displays the group activation for each condition.

Marmoset vocalizations (Figure 1B) elicited bilateral brain activations in primary auditory cortex, including the core (primary area [A1] and rostral field [R], rostral temporal [RT]), belt (caudomedial [CM], caudolateral [CL], mediolateral [ML], rostromedial [RM], anterolateral [AL], rostrotemporal medial [RTM], rostrotemporal lateral [RTL]), and parabelt areas (caudal parabelt [CPB], rostral parabelt [RPB]). Additionally, activations were found in V1, V2, GI, TE1, TH, the medial superior temporal area (MST), the temporo-parietal-occipital area (TPO), the superior temporal rostral area (STR), the retroinsular area (ReI), as well as in bilateral frontal areas, including primary motor cortex 4ab, somatosensory cortex areas 3a, 3b and 1/2, premotor areas 6DR, 6DC, cingulate area 24c and in left cingulate areas 32, 32v and 25. In the right hemisphere there were also activations in S2I, DI, Ipro, agranular insular cortex (AI), medial part of parainsular cortex (PaIM) and 8aV.

Scrambled vocalizations (Figure 1E) elicited responses mostly confined to the auditory cortex, with bilateral activations in core (A1, R, RT), belt (CM, CL, ML, RM, AL, RTM, RTL) and parabelt (CPB) cortices, as well as activations in right hemisphere in adjacent areas TPO, MST, STR, GI, GI, DI, S2I, Ipro and PaIM.

### Functional brain activations during the processing of audiovisual signals

In the intact audiovisual condition illustrated in Fig. 1C – where marmoset faces were combined with corresponding vocalizations – we identified activations that reflected the combination of the previously described unimodal maps (Figure 1A and 1B), alongside additional parietal, cingulate and frontal areas (Figure 1C). Specifically, we observed activations in the previously mentioned areas along the occipital-temporal axis (i.e., bilateral V1, V2, V2, V4, V4t, MT, 19DI, TEO, MST, FST, Pga-IPa, TPO, TE3, TE2, TE1, STR, 36, 35, Ent), in auditory regions of the core, belt and parabelt cortices (i.e., bilateral A1, R, RT, CM, CL, ML, RM, AL, RTM, RTL, RPB, CPB), as well as in regions of the prefrontal and premotor cortices (i.e., bilateral 8aD, 8Av, 6Va, 45, 47M, 47O and 6DR). Beyond these areas, we found activations in bilateral posterior parietal areas surrounding the intraparietal sulcus (IPS), including the occipito-pareital transitional areas (OPt), and the anterior, lateral, medial and ventral intraparietal areas (AIP, LIP, MIP and VIP), as well as PG, PFG, PE and PGM areas. Within the cingulate cortex, activations were present in bilateral areas 32, 25, 24a, 23b, 30 and 29d.

In the scrambled audiovisual condition depicted in Figure 1F – where scrambled marmoset faces were associated with scrambled corresponding vocalizations – there were strong activations in bilateral auditory areas (i.e., A1, R, RT, CM, CL, ML, RM, AL, RTM, RTL, RPB, CPB), and in adjacent areas STR, TPO, MST, ReI, FST and Pga-IPa. Temporal areas TE3 and TEO, visual areas V1, V2, V3, V4t, MT, V6, as well as parietal areas LIP, PFG and PG were also recruited. In the right hemisphere, we also observed activations in areas 8Av, 45, and in the insular areas GI, DI, AI.

When comparing marmoset faces paired with corresponding vocalizations to their scrambled versions (Figure 2) (i.e., marmoset face videos with corresponding vocalizations condition > scrambled marmoset face videos with corresponding scrambled vocalizations condition), we found stronger activations for the intact stimuli in the occipitotemporal, frontal and orbitofrontal cortices in bilateral areas V1, V2, V3, V4, MT, V4t, FST, Pga-IPa, TPO, TE3, TE2, TE1, 35, 36, 8Av, 6Va, 45, 47M, 13L, 47O, OPAI, as well as in left areas 8aD, 6DR, 13M and in right areas 8C and PaIM. However, no greater activations were found in the primary auditory cortex. Subcortically, greater activations were found in SC, pulvinar and amygdala (Figure 2). Our results indicate that the integration of audiovisual signals involves a broad network, which includes not only the distinct face and vocal processing networks but also extends to encompass parietal, cingulate, and prefrontal regions. Notably, the intact and coherent pairing of marmoset faces with vocalizations—compared to their scrambled and incoherent counterparts—showed preferential processing along the occipitotemporal axis, in both the lateral and medial prefrontal cortices, and within the anterior cingulate cortex, especially area 32.

**Figure 2.**
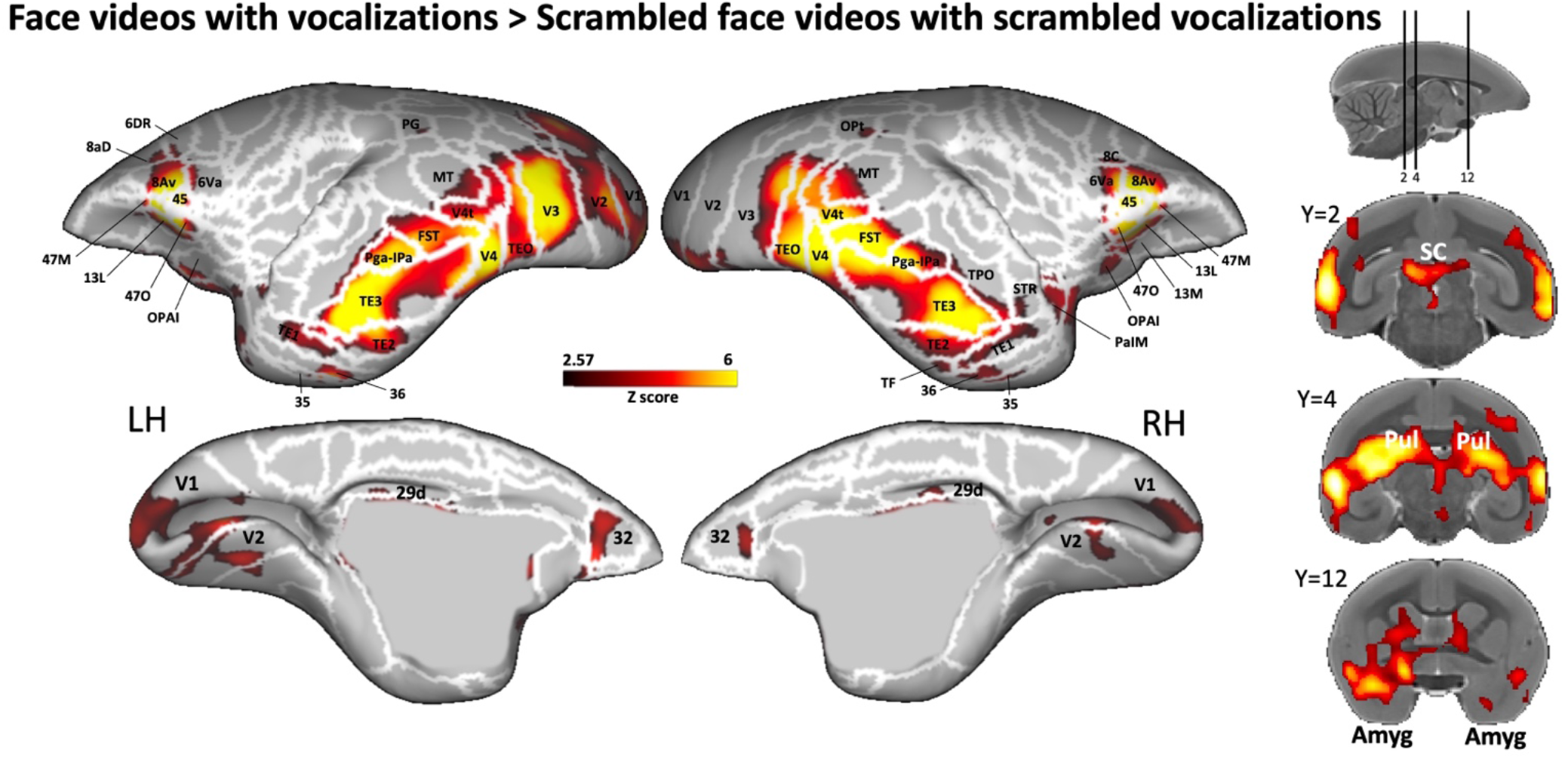
Brain network involved in processing intact *versus* scrambled conditions of combined face and vocal signals. Group functional maps illustrate significantly greater activations for the comparison between marmoset face videos paired with corresponding vocalizations and their scrambled versions. The group functional topology comparison is displayed on both the left and right fiducial marmoset cortical surfaces (lateral and medial views), as well as on coronal slices, to emphasize activations in subcortical areas. Regions are delineated by white lines, according to the Paxinos parcellation of the NIH marmoset brain atlas^86^. Reported brain areas have an activation threshold corresponding to z-scores > 2.57 (p<0.01, AFNI’s 3dttest++, cluster-size correction α=0.05 from 10000 Monte-Carlo simulations).

### Common and distinct brain regions involved in processing visual, auditory and audiovisual signals

To identify common and distinct brain regions engaged in processing visual, auditory, and audiovisual modalities, we conducted a conjunction analysis separately for intact and scrambled conditions. The results of this analysis are displayed in Figure 3A (intact stimuli) and Figure 3B (scrambled stimuli). Our findings revealed a substantial overlap between the visual and auditory activation maps with the multisensory map (depicted in yellow and purple in Figure 3).

**Figure 3.**
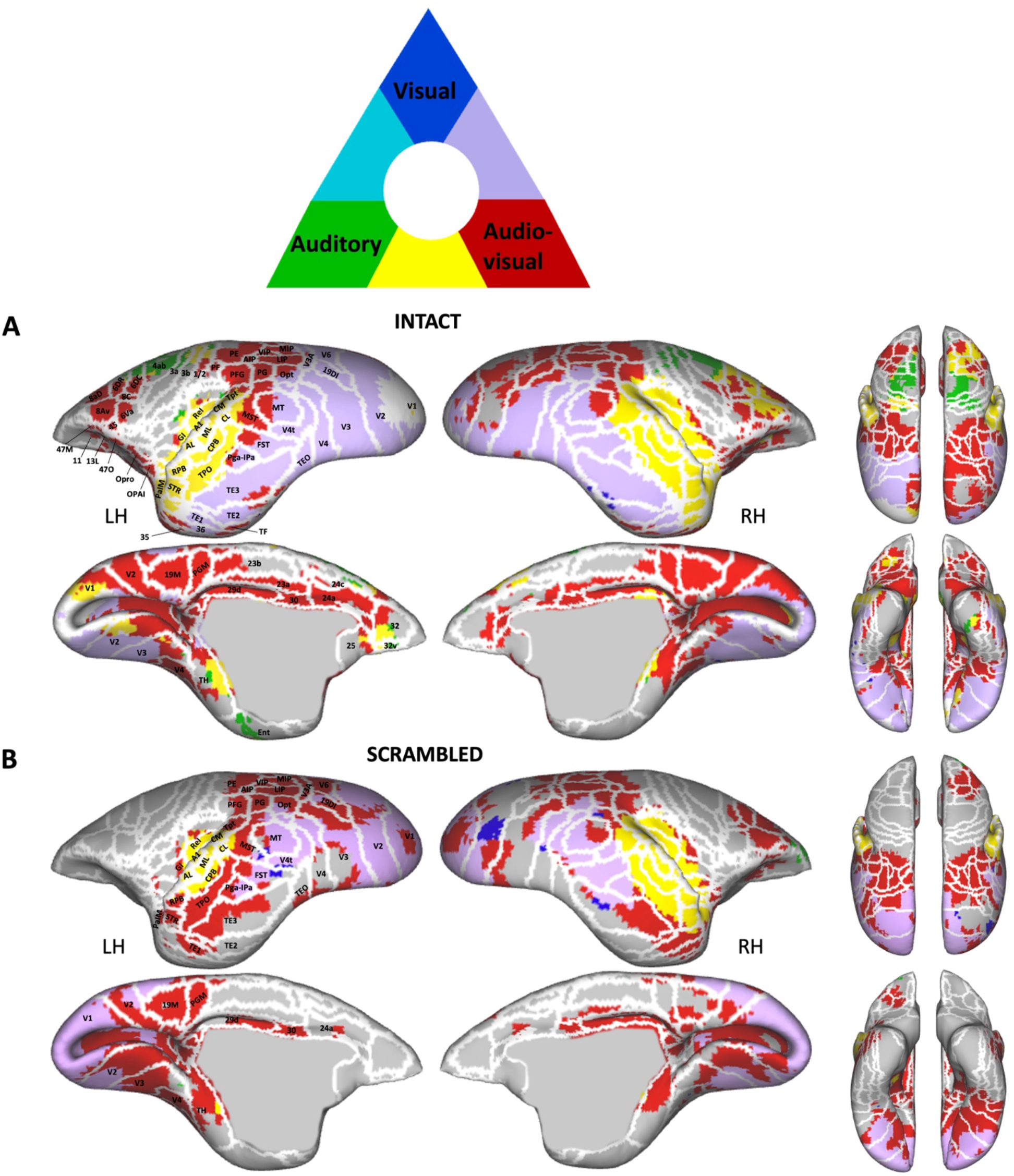
Spatial overlap of cluster networks for visual, auditory, and audiovisual processing in intact (A) and scrambled (B) conditions. Marmoset cortical surfaces of both hemispheres are shown, displaying all significant voxels (z-scores > 2.57; p<0.01, AFNI’s 3dttest++, cluster-size correction α=0.05 from 10000 Monte-Carlo simulations) as blue (unimodal visual condition > baseline), green (unimodal auditory condition > baseline), or red (audiovisual condition > baseline). Network overlap is indicated by the color key above the surface maps. Regions are delineated by white lines, according to the Paxinos parcellation of the NIH marmoset brain atlas ^86^.

Significantly, additional brain regions in the frontal, cingulate, and parietal cortices were found to be engaged only during multisensory stimulation (shown in red in Figure 3). This observation suggests that multisensory processing is not a simple summation of each modality but also involves additional brain regions. Both the intact and scrambled maps selectively activated bilateral posterior parietal areas (i.e., AIP, LIP, MIP, VIP, PG, PFG, PE, PGM) and cingulate areas (i.e., 29d, 30, 24a), with the intact condition further recruiting other cingulate areas (23b, 23a, 24c) and portions of rostral cingulate areas 25 and 32. Interestingly, the anterior portion of area 32 was solely activated by auditory stimulation (shown in green in Figure 3A), while its most posterior portion was only activated by multisensory stimulation (depicted in red in Figure 3A). Between these, the region was commonly activated by both auditory and multisensory conditions (indicated in yellow in Figure 3A).

In addition, certain prefrontal areas (8Av, 8aD) and premotor areas (6DR, 6DC) in the right hemisphere were shared between the auditory and multisensory intact maps (shown in yellow in Figure 3A). However, other bilateral prefrontal (45, 47M, 47O) and orbitofrontal areas (11, 13L, Opro, OPAI) were specifically activated by the intact multisensory condition (depicted in red in Figure 3A).

For the scrambled condition, only portions of areas 6DR, 6DC, 8C, 8Av, 8aD, 47M, 47O in the right hemisphere were activated by the audiovisual stimuli (Figure 3B, shown in red).

Overall, for the intact audiovisual conditions, regions responding primarily to visual stimulation (and not auditory) were predominantly concentrated in the occipital and temporal cortices (Figure 1A). In contrast, areas activated by auditory stimulation (but not visual) were situated in the primary auditory cortex, premotor cortex, and inferior frontal cortex (Figure 1B). These regions also responded to combined audiovisual stimulation (Figure 1C and 3A, shown in yellow and purple). Notably, no regions were identified that responded to both unimodal auditory and visual conditions (absence of light blue in Figure 3A). Areas responsive to combined audiovisual stimulation, but not to unimodal visual and auditory stimulation, were distributed across frontal, cingulate, and parietal regions (Figure 3A, shown in red). For the scrambled audiovisual conditions, regions responding to visual but not auditory stimuli remained concentrated in the occipital and temporal cortex, albeit to a lesser extent (Figure 1D). The areas activated by auditory but not visual stimuli were primarily situated in the primary auditory cortex (Figure 1E). These regions also responded to combined audiovisual stimulation (Figure 1F and 3B, shown in yellow and purple). In this scrambled condition, areas solely responsive to audiovisual stimuli were mainly observed in parietal regions, with only a few in cingulate and right prefrontal regions (Figure 3B, shown in red).

In summary, the intact, coherent conditions engage a more expansive neural network than the scrambled, incoherent ones, underscoring the intricate interplay of marmoset faces and vocalizations within the brain. The integration of these specific cues thus seems to engage extra brain regions, beyond those commonly activated by general audiovisual multisensory stimuli, as evidenced in the scrambled multimodal scenario.

### Positive interaction for combined audiovisual signals: superadditive effect

To determine the superadditive effect – corresponding to an increased activation when subjects integrate multimodal information compared to the sum of activation from single-modality inputs – we created an activation map displaying regions with stronger activations for the combined auditory and visual face-related information conditions compared to the sum of responses from unimodal face and vocal conditions (i.e., face videos with corresponding vocalizations > face videos + vocalizations).

This contrast, as illustrated in Figure 4A for intact conditions and 4B for scrambled conditions, revealed that areas in the temporal (i.e., bilateral areas TEO, FST, MST, PGa-IPa, TE3, TE2, TE1, TPO, 36), parietal (i.e., bilateral areas MIP, LIP, VIP, AIP, PG, PE, PFG, PGM), cingulate (i.e., bilateral areas 23b, 23a, 29d, 30, 24a), as well as premotor and prefrontal (i.e., areas 8Av, 6va, 8C, 8aD, 6DR, 47M, 47O bilateral for intact and right for scrambled) cortices responded more robustly to combined audiovisual conditions compared to the cumulative response of isolated auditory and visual conditions. Furthermore, while visual (i.e., bilateral areas V1, V2, V3, V4, MT, V6, V3A, V4t, 19DI) and auditory (comprising bilateral regions like A1, CM, ML, CL, CPB, RBP) cortices also displayed notable responses differences between multimodal and unimodal stimulations, the most significant variance in activations was primarily located in the temporal, parietal, and cingulate regions.

**Figure 4.**
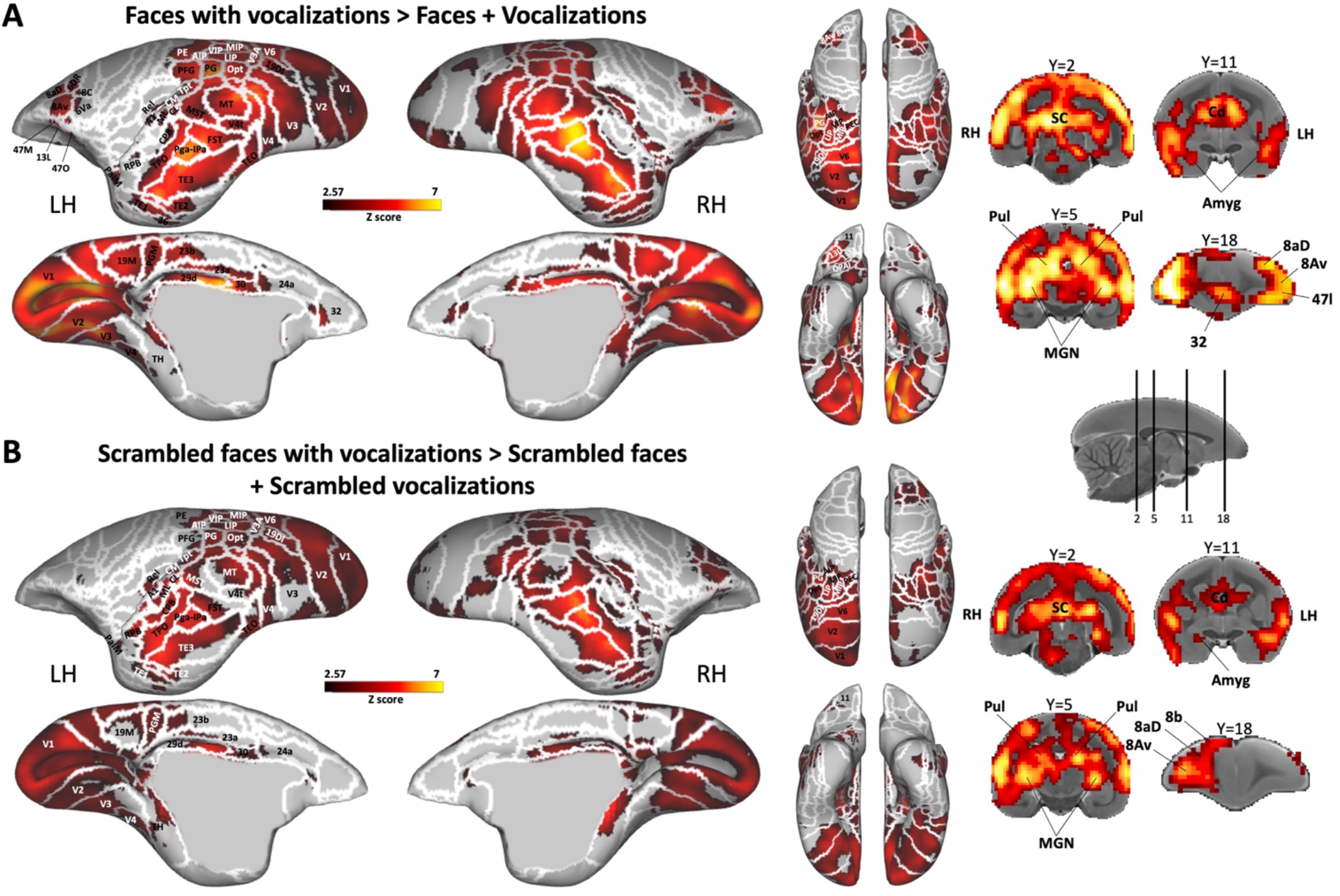
Superadditive neural processing of multisensory face and vocal signals. Group functional maps illustrate significantly greater responses to the multisensory audiovisual conditions compared to the sum of the responses for its unimodal constituents for both intact (A) and scrambled (B) stimuli. Significant differences were determined with paired t-tests, thresholded at z > 2.57 (p<0.01, AFNI’s 3dttest++, cluster-size correction α=0.05 from 10000 Monte-Carlo simulations). The group functional topology comparisons are displayed on both left and right fiducial marmoset cortical surfaces, as well as on coronal slices. Regions are delineated by white lines, according to the Paxinos parcellation of the NIH marmoset brain atlas ^86^.

Subcortically, structures such as the superior colliculus, caudate, MGN, pulvinar, and amygdala exhibited a stronger response to the multisensory condition compared to the combined responses of unimodal conditions.

Overall, this multimodal enhancement effect was more pronounced and extensive for intact conditions compared to scrambled conditions. Notably, area 32 was activated solely by the intact contrast.

## Discussion

Our study aimed to identify the distinct and shared neural substrates responsible for processing marmoset faces, vocalizations, and the combination of marmoset faces with their associated vocalizations. Within this multisensory framework, we further examined how these integrated audiovisual signals are processed in comparison to unimodal auditory and visual stimuli. The ability to recognize and integrate auditory and visual cues is crucial for effective communication. Our central hypothesis proposed that marmosets process face and vocal signals in distinct face and vocal ‘patches’ within their temporal and frontal cortices, and that these ‘patches’ should also be involved in audiovisual multisensory processing, with responses to combined stimuli exceeding the summed responses of individual auditory and visual stimuli – a phenomenon known as superadditivity.

To test this hypothesis, we utilized ultra-high field fMRI acquisitions and exposed awake marmosets to various stimuli, including marmoset face videos, vocalizations, and their corresponding scrambled versions both separately and in combination.

### Face and Vocalization Processing: Face-patch and Voice-patch systems

The ability to recognize faces is paramount in deciphering the intentions of others, making the differentiation and interpretation of facial expressions vital for social communication ^55–57^. Previous fMRI research on face processing in human and nonhuman primate has identified ‘face patches’ located across temporal and prefrontal cortices, which responded strongly to faces compared to non-face objects or scrambled faces ^6–8,10,11,16–19^. Our results for visual stimulation depicting marmoset face videos are consistent with previous studies and support recent investigations into face processing in marmosets, emphasizing the role of occipito-temporal regions in processing faces and facial expressions ^16–18^. These areas display robust activation in areas V2/V3, V4/TEO, V4t/FST, TE2-TE3, corresponding to the previously identified face-patches in marmosets (i.e., patches O (occipital), PV (posterior ventral), PD (posterior dorsal), MD (middle dorsal) and AD (anterior dorsal) respectively) ^16,18^.

Research in primate vocalization processing, another fundamental mode of communication in primates, has revealed the existence of selective ‘voice patches’ in the temporal cortex, specifically along the superior temporal sulcus (STS), and in certain regions of the frontal cortex in humans, macaques, and marmosets. These regions preferentially responded to vocalizations over other sounds or scrambled vocalizations ^29–32,36–39,42,58^. Our results align with these findings and our recent investigation in marmosets ^42^, showing similar activations in the primary auditory cortex, rostral cingulate, frontal, and temporal cortices for auditory stimulation playing vocalizations. We also confirmed similar primary auditory cortex activations previously obtained for the scrambled vocalizations.

### Multisensory Integration of face and vocal signals: involvement of temporal, parietal, prefrontal and cingulate cortices

Multisensory integration is defined as a process whereby the neuronal response to two sensory inputs is different from the sum of the neuronal responses to each on its own ^59,60^. Numerous fMRI studies in humans and several electrophysiological studies in macaques have demonstrated audiovisual multisensory interactions in the temporal cortex. Specifically, the STS has been identified as displaying multisensory responses in both humans and macaques, exhibiting enhanced activity for bimodal auditory and visual signals over unimodal ones. Human fMRI studies have confirmed the STS as a multisensory region ^51,52^, emphasizing its specialization in integrating various types of information within modalities (e.g., visual form, visual motion) and across modalities (auditory and visual). In macaques, electrophysiological evidence has revealed that individual neurons in the STS may respond solely to auditory stimuli, exclusively to visual stimuli, or to both auditory and visual stimuli ^61,62^. Additionally, some research has indicated that STS also harbors cells with selective audiovisual responses to faces ^46,63,64^.

Beyond the STS, recent evidence of audiovisual integration during naturalistic social stimuli has been found in specific regions of the monkey face-patch system ^47^, the voice-patch system ^44–46^, and the prefrontal cortex with the presence of multisensory neurons in the ventrolateral prefrontal cortex (VLPFC) responsive to combined face and voice stimuli ^48–50^.

Understanding the bimodal integration of visual and auditory signals in primate has previously been derived exclusively from human and macaque studies, leaving the processing of associations between vocalizations and dynamic faces in marmosets unexplored until now.

In our study, the multisensory condition, which combined marmoset face videos and corresponding vocalizations, increased the activity of face and voice ‘patches’, illustrating the similarity of these areas’ role in audiovisual multisensory integration in marmosets with what has been observed in macaques. Moreover, the activations seen in the TE complex (i.e., TE3, TE2, and TE1) could correspond to the STS responses observed in macaques and humans (Yovel and Freiwald, 2013), further showcasing multisensory integration in these regions in marmosets. A recent electrophysiology study in macaques established that neurons in the anterior fundus face patch in the STS (patch AF) responded to both visual and vocal stimuli, signifying their role in audiovisual integration during social communication ^47^. Our findings support this, showing activations in the temporal ‘face-patches’ regions during the integration of faces and vocalizations. The activations observed in the anterior face patches MD and AD might be equivalent to the AF patch in macaques ^11^.

In the frontal cortex, we also observed activations in various prefrontal areas, unveiling their multisensory role during processing of simultaneous facial expressions and vocalizations. Significantly, the rostral cingulate area 32, previously found to be activated in response to vocalizations in marmosets ^42^ and during language processing in humans ^31,65,66^, was also engaged in voice-face multisensory processing in our study.

Our conjunction analysis revealed that the processing of audiovisual stimulation extended beyond the recruitment of areas solely responsible for unimodal visual and auditory processing. This included not only the regions that process face and vocal signals but also additional cortical areas, suggesting a more intricate network engaged in multisensory integration. This finding highlights the complexity of multisensory processing, shedding new light on the underlying neural mechanisms. The areas activated solely by the multisensory condition were primarily situated in parietal, frontal and cingulate cortices, indicating that these regions may also serve as integrative hubs necessary to process the integration of both visual and auditory information for more efficient social communication.

Firstly, our findings reveal a noteworthy pattern of bilateral posterior parietal activations in response to combined audiovisual stimuli, around the IPS and in parietal areas PF, PE, PFG and PGM. This finding aligns with and extends existing research conducted on humans and macaques, which implicates parietal areas in multisensory processing ^67,68^. In humans, numerous fMRI studies have revealed the involvement of the IPS in tasks that demand the integration of visual and auditory stimuli ^53^, aligning well with our present observations in marmosets. In macaques, although some studies suggest audiovisual integration in the posterior parietal cortex, responses to stimuli in bimodal conditions have not been directly examined ^69^. Our results with marmosets parallel the findings observed in humans, underscoring the potential evolutionary conservation of this region’s role in multisensory processing across primates.

Secondly, our results reveal several areas in the frontal cortex situated in some lateral prefrontal (i.e., 47M, 47O, 45) and orbitofrontal areas (i.e., 11, 13L, OPro, OPAI), as well as in posterior and anterior cingulate areas (i.e., 29d, 23d, 23a, 30 24a) responding only to the multisensory audiovisual condition. In macaques, the integration of auditory and visual information has been described in the VLPFC. Some studies have shown that single neurons in VLPFC integrate audiovisual species-species face and vocal communication stimuli, suggesting that these neurons are an essential node in the cortical network composed by unimodal auditory and visual regions responsible for communication ^48–50^.

In humans, fMRI studies have also demonstrated the activation of inferior frontal gyrus during the processing and integration of speech and gestures ^70,71^, suggesting a larger role of the IFG in communication than classical auditory-speech processing ^72^. Some studies have further demonstrated a decrease in ventral prefrontal activity for incongruent faces and voices ^60,71^. In macaque monkey, the evidence shows that cells in the ventral PFC respond to and integrate audiovisual information, with some cells exhibiting multisensory enhancement or suppression when face-vocalization stimuli are combined ^48,49^. In humans, the integration of audiovisual stimuli also occurred at the level of the anterior cingulate/medial prefrontal cortex ^54^. It has been shown that the activity in these areas was enhanced during pairing of congruent and incongruent cross-modal visual and auditory stimuli, and the activation was greater during matching conditions. These results align with our findings in marmosets, with activations for the combined audiovisual conditions in cingulate/medial prefrontal cortex. Thus, our results on marmosets have allowed us to add a piece of evidence that the ventral frontal lobe and the cingulate cortex of primates may be involved in processing the association between a face or facial gesture and a vocal stimulus. This suggest that the PFC may be a precursor to the more complex functions of the human frontal lobe, where semantic meaning is linked with acoustic or visual symbols ^49^. Future studies in marmosets would be necessary to determine if, like humans^54^, these activations would be even more pronounced in congruent compared to incongruent pairing of these face and vocal signals.

It is important to note that, although not previously reported in macaques and humans, we identified that the rostral cingulate area 32 participates in the integration of visual and auditory stimuli. Interestingly, only the posterior portion of this areas responded exclusively to multisensory stimulation. In contrast, the anterior portion activated solely in response to auditory cues. The intervening region displayed activation in both auditory and combined audiovisual conditions. Nevertheless, these findings held true only for intact, coherent stimuli and not for the scrambled, incoherent versions. While the neural network for multisensory processing between intact and scrambled conditions bear resemblance, the activations were less strong and less extended for the combined scrambled audiovisual stimuli compared to their intact counterparts. This disparity was especially pronounced in the prefrontal cortex and was less evident in the parietal cortex. Directly comparing the two conditions, intact audiovisual stimuli elicited greater activations across the occipito-temporal axis, in the prefrontal cortex, and in the rostral cingulate area 32. Subcortically, the superior colliculus, pulvinar and amygdala were also more activated by the intact audiovisual condition. The activation patterns in the rostral cingulate area 32, especially the distinct responses between its posterior and anterior portions, suggest a specialized role in the integration of coherent and congruent multisensory information. Its selective response to intact stimuli over scrambled stimuli, emphasize its pivotal role in processing structured and meaningful combinations of audiovisual social cues. This region’s involvement in processing congruent multisensory signals, particularly within the context of social interactions, may be crucial for interpreting and responding to complex social stimuli. The differential activation across the temporal, prefrontal and cingulate regions between intact and scrambled conditions could represent a sophisticated neural mechanism that distinguishes relevant audiovisual social information from nonsensical or mismatched stimuli, thereby enhancing the precision and efficacy of primate social communication.

The principle of superadditivity, in which multisensory responses exceed the sum of the linear additive responses to the unimodal stimuli, has been advocated by some researchers as a requirement for brain regions involved in multisensory integration ^73,74^. In line with this, we assessed the augmented responses to the audiovisual multimodal conditions in comparison to the sum of visual and auditory unimodal conditions, aiming to identify the regions exhibiting a superadditive effect. Our findings revealed that this augmented response was not merely a simple summation of unimodal stimuli, but rather a complex interplay of activation across temporal, parietal, cingulate, lateral, and medial prefrontal areas. This suggests a synergistic enhancement of sensory processing that goes beyond mere additive effects. Both visual and auditory areas also demonstrated significant differences between multimodal and unimodal stimulations, although the more pronounced differences in activations were situated in the temporal, parietal, and cingulate areas. Subcortically, structures such as the superior colliculus, caudate, MGN, pulvinar, and amygdala exhibited a stronger response to the multisensory condition compared to the sum of unimodal conditions. As previously described, we also found pronounced and extensive multimodal enhancement effects for intact, coherent conditions as opposed to scrambled, incoherent conditions. Notably, some lateral prefrontal areas and the rostral cingulate area 32 were activated solely by the intact contrast. This phenomenon may be instrumental in enriching the perception of stimuli that are congruent across different sensory modalities, thereby facilitating a more coherent and integrated representation of the external environment. The superadditive effect in these regions likely contributes to the efficiency and robustness of multisensory integration, enhancing the marmosets’ ability to perceive, interpret, and respond to complex social cues. These findings not only confirm the concept of superadditivity in the neural processing of audiovisual stimuli but also offer new insights into the functional organization and adaptability of the neural networks underlying social communication in primates.

In summary, our findings suggest that the face ‘patches’ along the occipitotemporal axis, vocal ‘patches’ in the auditory cortex, and their interconnections with prefrontal and cingulate areas potentially constitute a conserved audiovisual multisensory pathway in marmosets. This indicates that, during audiovisual social communication, a more expansive network is at play than when processing visual and auditory signals individually. Considering the highly social nature of marmosets, these findings underscore the importance of concurrently integrating facial and vocal cues, thereby helping marmosets in decoding and interpreting social signals during their interactions ^4,75–79^.

## Methods

### Common Marmoset Subjects

All experimental procedures were in accordance with the guidelines of the Canadian Council of Animal Care policy and a protocol approved by the Animal Care Committee of the University of Western Ontario Council on Animal Care #2021-111. Ultra-high field fMRI data were collected from six awake common marmoset monkeys (*Callithrix jacchus*): two females (weight 315 and 150 g, age 44 months) and four males (weight 365-459 g, age 32-44 months).

To prevent head motion during MRI acquisition, animals were surgically implanted with an MR-compatible machined PEEK (polyetheretherketone) head post ^80^, conducted under anesthesia and aseptic conditions. During the surgical procedure (for details, see ^81^, the animals were first sedated and intubated to ensure they remained under gas anesthesia, maintained by a mixture of O2 and isoflurane (0.5-3%). With their heads immobilized in a stereotactic apparatus, a machined PEEK head fixation post was positioned on the skull following a midline skin incision along the skull. This device was secured in place using a resin composite (Core-Flo DC Lite; Bisco). Heart rate, oxygen saturation, and body temperature were continuously monitored throughout the surgery. Two weeks post-surgery, the monkeys were acclimated to the head-fixation system and the MRI environment through a three-week training period in a mock scanner, as described in Gilbert et al., (2021).

### Multisensory facial expressions task and experimental setup

We utilized six different types of stimuli derived from previously recorded videos ^18^. These videos originally depicted negative facial expressions. From these videos, we generated two video conditions, two audio conditions and two conditions involving both video and corresponding audio (Figure 5B). We used custom video-editing software (iMovie, Apple Incorporated, CA) for this purpose. The two video conditions encompassed marmoset face videos with no sound and their scrambled versions. The two audio conditions consisted of vocalizations extracted from the videos and their scrambled counterparts. The final two conditions combined the videos and corresponding audio, and their scrambled versions. No vocalization filtering or background noise cancellation was implemented to maintain the integrity of the vocal features.

**Figure 5.**
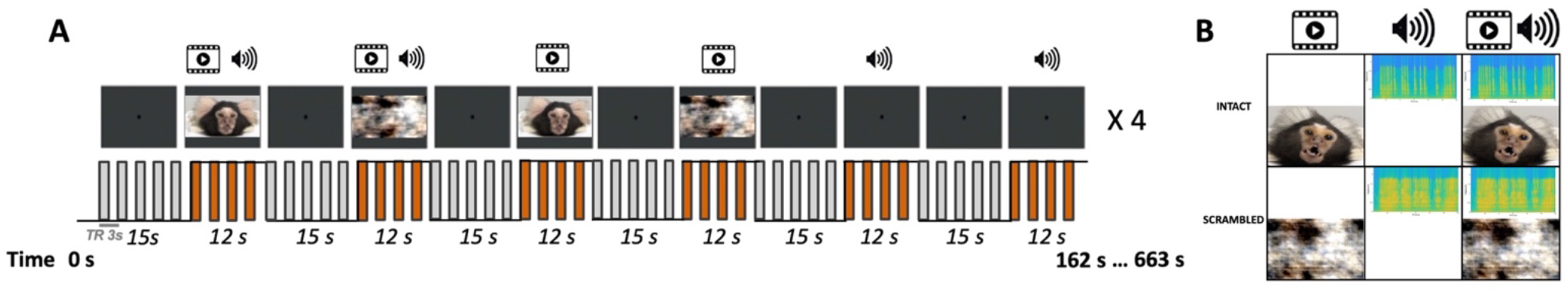
Experimental setup and stimuli. A. A sparse fMRI block design was employed, featuring a repetition time (TR) of 3 seconds. Each TR was preceded by a silent period of 1.5 seconds to ensure proper perception of stimuli. During each run, six different conditions were presented in a randomized order, each lasting for 12 seconds and repeated four times. Overall, there were 24 stimulus blocks interleaved with 23 baseline blocks lasting 15 sec, during which a central dot was displayed in the center of the screen. **B**. The conditions comprised three intact and three scrambled conditions: 1) a unimodal condition featuring video of marmoset faces, 2) a unimodal condition of marmoset vocalizations, and 3) a multimodal condition displaying marmoset faces with corresponding vocalizations.

As in our previous study ^18^, we phase-scrambled the videos with a custom program, while vocalizations were time-domain scrambled ^83^. This preserved their spectral content over longer time periods but removed structure at shorter timescales, rendering the vocalizations unintelligible ^42^.

Each condition was incorporated in a block design task, wherein each block of stimuli lasted twelve second, interleaved with a fifteen-second baseline block. During the baseline block, a 0.36° circular black cue was displayed at the screen center against a gray background. Each run repeated the six conditions four times. To randomize the presentation of conditions in each run, we created eight different stimulus sets. These were counterbalanced within and between animals (Figure 5A).

During the scanning sessions, monkeys were placed in a horizontal MR scanner (9.4 T) in a sphinx position within an MRI-compatible restraint system. Their heads were secured using a head post, and MRI-compatible auditory tubes were worn ^42,80^. After performing the head fixation steps, the MRI-compatible auditory tubes (S14, Sensimetrics, Gloucester, MA) were directly placed into the animals’ ear canals bilaterally and were fixed using reusable sound-attenuating silicone earplugs (Amazon) and self-adhesive veterinary bandage.

An MR-compatible camera (model 12M-i, MRC Systems GmbH, Heidelberg, Germany) was positioned to monitor the animal during acquisition. Horizontal and vertical eye movements at were tracked at a frequency of 60Hz using a video eye tracker (ISCAN ETL-200 system, Boston, Massachusetts). Analysis of functional run data was performed using a custom R script. In each experimental condition and during baseline periods (i.e., fixation point in the center of the screen), the animals spent more than 79% of the time looking at the screen (baseline: 82.6%; marmoset face videos: 83.7%; vocalizations: 84.9%; marmoset face videos with vocalizations: 90.4%; scrambled marmoset face videos: 79.4%; scrambled vocalizations: 82.4%; scrambled marmoset face videos with corresponding scrambled vocalizations: 83.5%). Although a one-way ANOVA test indicated an effect of condition on viewing time (F(2.62,13.10) = 3.69, p = 0.044), this result was not robust after Bonferroni correction for multiple comparisons (minimum adjusted p-value = 0.58). Consequently, any differences in fMRI activation between the conditions cannot be attributed to differences in exposure to the stimuli.

Visual stimuli were projected onto a forward-facing plastic screen positioned 119 cm from the animal’s head using an LCSD-projector (Model VLP-FE40, Sony Corporation, Tokyo, Japan) via a back-reflection on a first surface mirror. We used Keynote software (version 12.0, Apple Incorporated, CA) for stimulus display. The onset of each stimulus was synchronized with an MRI TTL pulse triggered by a python program running on a Raspberry Pi (model 3B+, Raspberry Pi Foundation, Cambridge, UK). The animals received a reward only before and after each scanning session, not during sessions.

### fMRI acquisition and parameters

Imaging was performed at the Center for Functional and Metabolic Mapping at the University of Western Ontario. Data was collected using a 9.4T/31 cm horizontal bore magnet and a Bruker BioSpec Avance III console running the Paravision-7 software package (Bruker BioSpin Corp). A custom-built gradient coil, with a 15-cm inner diameter and maximum gradient strength of 1.5 mT/m/A, coupled with eight separate receive channels was employed ^80^.

Eight functional images were acquired per animal over varying sessions, depending on each animal’s compliance. We utilized a gradient-echo based single-shot echo-planar images (EPI) sequence, with parameters set as follows: TR=3s, acquisition time TA=1.5s, TE = 15ms, flip angle = 40°, field of view=64×48 mm, matrix size = 96×128, isotropic resolution of 0.5 mm3, 42 axial slices, bandwidth=400 kHz, and a GRAPPA acceleration factor of 2 (left-right). An additional set of EPIs, featuring an opposite phase-encoding direction (right-left), was collected for the EPI-distortion correction.

To diminish potential auditory stimuli masking by scanner noise, we employed a continuous acquisition paradigm that incorporated silent periods ^42^. Despite the continuous presentation of auditory stimuli during each 12-second stimuli block, 1.5-second periods within each 3-second TR experienced the scanner noise level turning off. Consequently, we used a 3-second TR but gathered all slices within 1.5 seconds.

In each session per animal, we also acquired a T2-weighted structural image, with parameters set as follows: TR=7s, TE=52ms, field of view=51.2×51.2 mm, resolution of 0.133×0.133×0.5 mm, 45 axial slices, bandwidth=50 kHz, and a GRAPPA acceleration factor of 2.

### fMRI preprocessing

The data was processed using AFNI ^84^ and FMRIB/FSL ^85^ software packages. Initially, the raw functional images were converted into NifTI format using AFNI’s dcm2nixx function, and then reoriented from the sphinx position using FSL’s fslswapdim and fslorient functions. The functional images were despiked with AFNI’s 3Ddespike function, and volume were registered to the middle volume of each time series using AFNI’s 3dvolreg function. We stored the motion parameters from volume registration for later use with nuisance regression. Subsequently, functional images were smoothed using a full width at half-maximum Gaussian kernel (FWHM) of 1.5mm with AFNI’s 3dmerge function and bandpass filtered from 0.1 to 0.01 Hz with AFNI’s 3dBandpass function. For each run, an average functional image was calculated and linearly registered to the respective T2-weighted anatomical image of each animal using FSL’s FLIRT function. For this process, the T2-weighted anatomical images were manually skull-stripped, and the mask of each animal was applied to the corresponding functional images. The transformation matrix obtained after the registration was used to transform the 4D time series data.

Lastly, the T2-weighted anatomical images were registered to the NIH marmoset brain atlas ^86^ via nonlinear registration using Advanced Normalization Tools (ANTs’ ApplyTransforms function).

### fMRI statistical analysis

We used AFNI’s 3dDeconvolve function to estimate the hemodynamic response function (HRF), for each run. The task timing was initially convolved with the time series data to estimate the HRF for each of the six conditions, using AFNI’s ‘BLOCK’ convolution. Then, the amplitude and timing of the HRF for each condition were estimated, leading to the creation of a regressor for each condition to be used in the subsequent regression analysis. Subsequently, all the conditions were entered into the same model, in conjunction with polynomial detrending regressors and motion parameters, allowing for the evaluation of the statistical significance of the estimated HRF. This regression produced six T-value maps per animal per run, each corresponding to our experimental conditions. These resultant regression coefficient maps were then registered to the NIH marmoset brain atlas template space ^86^ using the transformation matrices obtained with the registration of anatomical images on the template (see above).

These maps were then subject to group level comparison via paired t-tests using AFNI’s 3dttest++ function, resulting in Z-value maps. To protect against false positives and control for multiple comparisons, we applied a clustering method derived from 10,000 Monte Carlo simulations to the resultant z-test maps using the ClustSim option (α=0.05). This method involves setting a cluster-forming threshold of p<0.01 uncorrected, followed by applying a family-wise error (FWE) correction of p<0.05 at the cluster-level.

These Z-value maps were displayed on fiducial maps obtained from the Connectome Workbench (v1.5.0, ^87^) using the NIH marmoset brain template ^86^, and on coronal sections. The Paxinos parcellation of the NIH marmoset brain atlas ^86^ was used to define anatomical locations of cortical and subcortical regions.

Initially, we identified voxels that exhibited significantly stronger activation during task engagement in each condition by contrasting each condition with the baseline period (i.e., marmoset face videos > baseline, scrambled marmoset face videos > baseline, vocalizations > baseline, scrambled vocalizations > baseline, marmoset face videos with vocalizations > baseline and scrambled marmoset face videos with scrambled vocalizations > baseline). To identify the areas involved in the processing of each modality (i.e., video, audio, and audiovisual conditions) and those common between modalities, we created a conjunction map between the three conditions, separately considering both intact and scrambled conditions. We used the AFNI 3dcalc-step function, inputting the thresholded resultant z-test maps obtained through paired t-tests (as described above). This allowed us to observe specific activations for each condition and the shared activations between conditions. Finally, to identify multisensory brain regions more activated by intact compared to scrambled audiovisual stimuli, we contrasted the activations for marmoset face videos with vocalizations against their scrambled versions (i.e., marmoset face videos with vocalizations > scrambled marmoset face videos with scrambled vocalizations).

Subsequently, we investigated the superadditive effect to determine the voxels more activated by combined audiovisual stimulation compared to the summed responses of unimodal auditory and visual stimulation. This comparison involved contrasting the multisensory condition (i.e., faces with corresponding vocalizations) with the summed response of the unimodal conditions (i.e., face video conditions plus vocalization conditions).

## Acknowledgements

Support was provided by the Canadian Institutes of Health Research (FRN 148365), the Canada First Research Excellence Fund to BrainsCAN, and a Discovery grant by the Natural Sciences and Engineering Research Council of Canada to SE. We wish to thank Cheryl Vander Tuin, Whitney Froese, Hannah Pettypiece, and Miranda Bellyou for animal preparation and care, and Dr. Alex Li for scanning assistance.

